# Urban fox squirrels exhibit tolerance to humans but respond to stimuli from natural predators

**DOI:** 10.1101/2020.12.18.423423

**Authors:** Anna Kittendorf, Ben Dantzer

## Abstract

Animals in urban areas that experience frequent exposure to humans often behave differently than those in less urban areas, such as exhibiting less vigilance or anti-predator behavior. These behavioral shifts may be an adaptive response to urbanization, but it may be costly if animals in urban areas also exhibit reduced anti-predator behavior in the presence of natural predators. In trials with only a human observer as the stimulus, urban squirrels exhibited reduced vigilance and anti-predator behavior compared to those in less urban areas. Next, we exposed squirrels in multiple urban and less urban sites to acoustic playbacks of a control stimulus (non-predatory bird calls), a natural predator (hawk), and dogs and recorded their vigilance and three different anti-predator behaviors when a human approached them while either broadcasting one of these three playbacks or no playbacks at all. Squirrels at urban sites also did not differ in their behavioral responses to the playbacks from possible predators (hawks or dogs) when they were compared to those at less urban sites exposed to these playbacks. Urban squirrels also exhibited increased vigilance and anti-predator behavior when exposed to a human paired with hawk playbacks compared to the control playbacks. Together, our results indicate that urban squirrels did perceive and assess risk to the natural predator appropriately despite exhibiting increased tolerance to humans. These results provide little support for the hypothesis that increased tolerance to humans causes animals to lose their fear of natural predators.

## Introduction

Behavior plays an important role in enabling animals to persist through environmental change (Baldwin, 1896; Bartholomew, 1964; West Eberhard, 1989; Price et al., 2003; Snell-Rood, 2013). Accordingly, it seems to play a major role in facilitating the ability of animals to cope with new challenges that they face in urban environments (Ditchkoff et al., 2006; Tuomainen and Candolin, 2011; Lowry et al., 2013; Ryan and Partan, 2014). One of the most common behavioral adjustments of animals in urban environments is reduced anti-predator behavior in the presence of humans. This is often reflected in measures of flight initiation distance (FID), which is the distance at which an animal flees from an approaching human and is thought to be an approximation of their sensitivity to risk of an approaching predator (Cooke, 1980; Ydenberg & Dill, 1986; Lima and Dill, 1990). Individuals with shorter FIDs are considered to be bolder than individuals with longer FIDs since they demonstrate reduced fear of the “predator”.

Substantial evidence supports the hypothesis that animals in more urban environments exhibit less anti-predator behavior, as reflected by a lower FID. For example, a meta-analysis of 180 bird species, 16 lizard species, and 16 mammal species, Samia et al. (2015) showed that populations of these species that experienced elevated levels of human disturbance exhibited lower FID. This could be because vigilance and anti-predator behaviors carry energetic or time costs as they take time away for resource acquisition and animals in urban environments may therefore optimize resource acquisition by exhibiting lower levels of anti-predator behavior (Ydenberg and Dill, 1986; Cooper and Frederick, 2007; Møller, 2012).

Although reductions in the expression of anti-predator in urban environments is generally thought to be adaptive (Møller, 2008; Carrette et al., 2016), there may be potential costs for urban animals if they reduce their overall expression of anti-predator behavior to not only humans but also towards natural predators if those urban areas contain predators. This could be due to the phenomenon of cross-habituation or stimulus-generalization. For example, birds that are habituated to a threatening stimulus that are then presented with a second simulated predator exhibit an attenuated response to this second stimulus compared to a group of naïve birds (Hinde, 1954; see also Curio, 1993). This type of stimulus generalization can occur where an animal habituated to one stimulus exhibits an attenuated response to a second stimulus from the same *or* different sensory modality (Guttman and Kalish, 1956; Thompson and Spencer, 1966; Rankin et al., 2009). Related concepts occur in the context of “behavioral spillover” where individuals that exhibit high levels of a behavior in one context also exhibit it in another context even though it may not be adaptive, such as animals exhibiting higher levels of boldness in a courtship context also exhibiting higher boldness in the presence of a predator (Arnqvist and Henriksson, 1997; Sih et al., 2004).

If urban animals in areas containing predators exhibit reductions in vigilance and/or anti-predator behavior not only toward humans but also to natural predators, it could conceivably have important impacts on wildlife populations by increasing their vulnerability to predators (Geffroy et al., 2015). To date, there is little consensus about whether animals in urban areas or those exposed to increased human presence exhibit a reduced response to threats from natural predators (Fitzgerald and Stronza, 2016). For example, some studies show that individuals in areas with higher human activity exhibit less of a behavioral response when natural predators were observed visiting the area (Olson and Acevedo-Gutiérrez, 2017) or due to acoustic playbacks of a natural predator (McCleery, 2009). The latter suggests that animals experiencing frequent exposure to human activity exhibit reduced responses to other stimuli from natural predators. Other studies show that the response of animals in more urban areas to a stimulus from a natural predator is not attenuated compared to those in more rural locations (Labra and Leonard, 1999; Coleman et al., 2008; Seress et al., 2011; Bokony et al., 2012; Cavalli et al., 2016; Weaver et al., 2018; Vincze et al., 2019).

In this study, we characterized the vigilance and anti-predator behavior of fox squirrels (*Sciurus niger*) in urban and less urban areas to achieve the following two objectives. First, we conducted standard FID trials (with only stimuli from a human observer) to examine whether squirrels in urban areas showed reduced vigilance and anti-predator behavior towards a human observer compared to those in less urban areas. If squirrels in urban areas did exhibit reduced vigilance and anti-predator behavior, this would support the hypothesis that squirrels in our urban study populations were more tolerant of human presence, which would be consistent with numerous other studies (Samia et al., 2015). Squirrels were located in their natural habitat and we recorded the following four aspects of their vigilance and anti-predator behavior. First, we recorded the distance to which the observer could get to before they exhibited vigilance behavior towards the observer (“first alert distance” or FAD, similar to Fernández-Juricic and Schroeder, 2003; Blumstein et al., 2005). Second, we recorded how close the observer could get to them before they ran away (FID). Third, was the probability that the squirrel escaped by running up a tree. Fourth, the latency following the trial it took them to resume their typical behavior (foraging or traveling off tree). We interpreted vigilance behavior was reflected in FAD and that anti-predator behavior was composed of FID, probability of the squirrel escaping up a tree, and the latency to resume typical behavior following the trial. However, we note that it is likely that all four of these behaviors are quite similar in the sense that they measured anti-predator behavior and that the latency to resume typical behavior following the trial may be affected by motivational issues associated with nutritional state. Measuring all four of them can provide additional insight, such as examining whether squirrels in less urban areas are more alert to human presence than those in urban areas. Additionally, most studies on this topic are in birds and only measure FID. Measuring whether the squirrel escaped by running up a tree and how long the squirrel took to resume their typical behavior in addition to FID may provide greater insight into the behavioral differences between animals in urban or less urban areas.

Our second objective was to examine whether urban animals exhibit reduced behavioral responses to stimuli from natural predators when they are in the presence of humans. To do so, we quantified the four behaviors described above when fox squirrels in urban or less urban areas were presented with a human observer with a control acoustic playback (common non-threatening bird), a human observer paired with the playback of a natural predator (hawk), or a human observer paired with a playback of an invasive predator (dog). We predicted that squirrels in the urban areas but not those in the less urban areas would exhibit no change in vigilance and anti-predator behavior when they were exposed to the human+dog or human+hawk stimuli compared to the human+control playback. We also predicted that squirrels in the urban sites would exhibit less vigilance and anti-predator behavior when exposed to hawk or dog playbacks compared to those at the less urban sites that were exposed to the hawk or dog playbacks. These results would support the hypothesis that animals in urban environments exhibit less vigilance and anti-predator not only to humans but also when faced with natural predators.

## Materials and Methods

### Study species and sites

Fox squirrels are ubiquitous in urban and suburban environments in the midwestern United States (McCleery, 2008, 2009). Although arboreal tree squirrel species like fox squirrels are common in urban areas worldwide, they continue to experience predation from natural predators, although it may be rare compared to other sources of mortality (McCleery et al., 2008). Urban squirrels also likely experience predation from domestic cats and dogs (Koprowski, 1994; Wauters et al., 1997; Tumlison, 2012; Loss et al., 2013; Jokimäki et al., 2017).

We studied natural populations of adult fox squirrels from six sites in and around Ann Arbor, Michigan (Table S1 in Appendix). Sites were chosen based upon estimates of human population density (Center for International Earth Science Information Network, 2018) with urban sites having higher human density than less urban sites (see below and Table S1). Urban sites included Prospect Park as well as two locations on the University of Michigan’s (UM) main campus (North and Central Campus) that are ∼3-4 km away from one another. Prospect Park is near downtown Ypsilanti, Michigan and about 13 km away from UM main campus. Less urban sites included Nichols Arboretum, County Farm Park, and Saginaw Forest. Nichols Arboretum is located ∼1 km away UM main campus, County Farm Park is about ∼4.5 km away, and Saginaw Forest is ∼7 km away. At all research sites, dogs are allowed but hunting is not. Squirrels may be occasionally fed by humans at some of our study sites (e.g., Central Campus), but data were not systematically collected to assess feeding rates. Approval to conduct this research at each site was obtained from UM (Central Campus, Nichols Arboretum, North Campus, Saginaw Forest), Washtenaw County Parks & Recreation (County Farm Park), and the City of Ypsilanti (Prospect Park). All of our field procedures were non-invasive and involved behavioral observation or short-term exposure to playbacks of acoustic stimuli. All experiments followed the guidelines set by the Animal Behavior Society/Association for the Study of Animal Behaviour (Anonymous, 2012) and the US National Research Council and were approved by the UM Institutional Animal Care and Use Committee (protocol # PRO00009076). We note that squirrels used in the study may be STRANGE (*sensu* Webster and Rutz, 2020) in the sense that individual squirrels likely have different rearing histories (though they are unknown and none should have been reared in captivity) and that compliance to take part in the study was likely biased towards squirrels that did not immediately run away when approached by the human observer.

Given that increased exposure to humans may cause animals in urban areas to exhibit less anti-predator behavior towards them (McCleery, 2009; Rodriguez-Prieto et al., 2009; Vincze et al., 2016; Uchida et al., 2019), we focused on human presence as the major factor difference between our study sites (which should also reflect general urbanization). Sites were classified as “urban” based upon having a human population density >1000 persons per km^2^ whereas the less urban sites had anywhere from 25-250 persons per km^2^ (Saginaw) to 250-1000 persons per km^2^ (County Farm Park, Nichols Arboretum). To support these classifications, we estimated human and dog presence while we were visiting sites conducting our behavioral observations. We counted the total number of dogs (on or off leash) but only counted the number of humans up to 50. If human presence exceeded 50 people, then a rough estimate of 50, 75, or 100 was recorded. Human presence was recorded as 100 for all numbers estimated to be >100. We did not record the distance from the observer to other humans but just whether the human was visible. Although human and dog presence varied among the different sites (Table S1), the number of humans observed per hour of observation averaged over all the urban sites (mean ± SE = 7.23 ± 3.33 humans/hr) was higher than those observed averaged over all the less urban sites (0.51 ± 0.40: Mann-Whitney-Wilcoxon Test, W = 1, *p* = 0.1). We observed fewer dogs per hour at the urban sites (0.065 ± 0.06 dogs/hr) compared to the less urban sites (0.33 ± 0.10, Mann-Whitney-Wilcoxon Test, W = 9, *p* = 0.1). These differences in humans or dogs observer per hour were not significant but in general support our assumption that our urban sites likely experience greater exposure to humans.

We aged and sexed squirrels visually according to their size (small juvenile squirrels were excluded) and anatomy (males were identified by presence of testes), respectively. Similar to most studies that measure anti-predator using FID, trials were conducted on unmarked squirrels at each site. We located squirrels by walking around each site and trials were started when squirrels were observed. Focal individuals were selected randomly, however, only squirrels that were feeding or foraging on the ground were included in this experiment. Because we did not mark squirrels individually, it is possible that the same squirrel was observed on different days, although we visited different areas of each study site to try and reduce this possibility. It is unlikely that the same squirrel was observed multiple times on the same day, because after each trial was completed, the observer walked approximately 20 meters away from the previous location (in a continuous linear direction from where the first trial was conducted) and started a trial with a different squirrel. Additionally, sites were only revisited after at least three days since the previous visit to reduce the possibility of a squirrel becoming habituated to the trials should it be sampled again. Although we cannot address habituation in this study, if squirrels at these sites were habituating to our protocols, we would expect that their behavioral responses would decline with trial number or date when the trial was conducted. The fact that none of our behavioral variables were associated with date of when the trial was conducted (Tables 1-2) supports our assumption that squirrels were not habituating to our protocols.

**Table 1.**
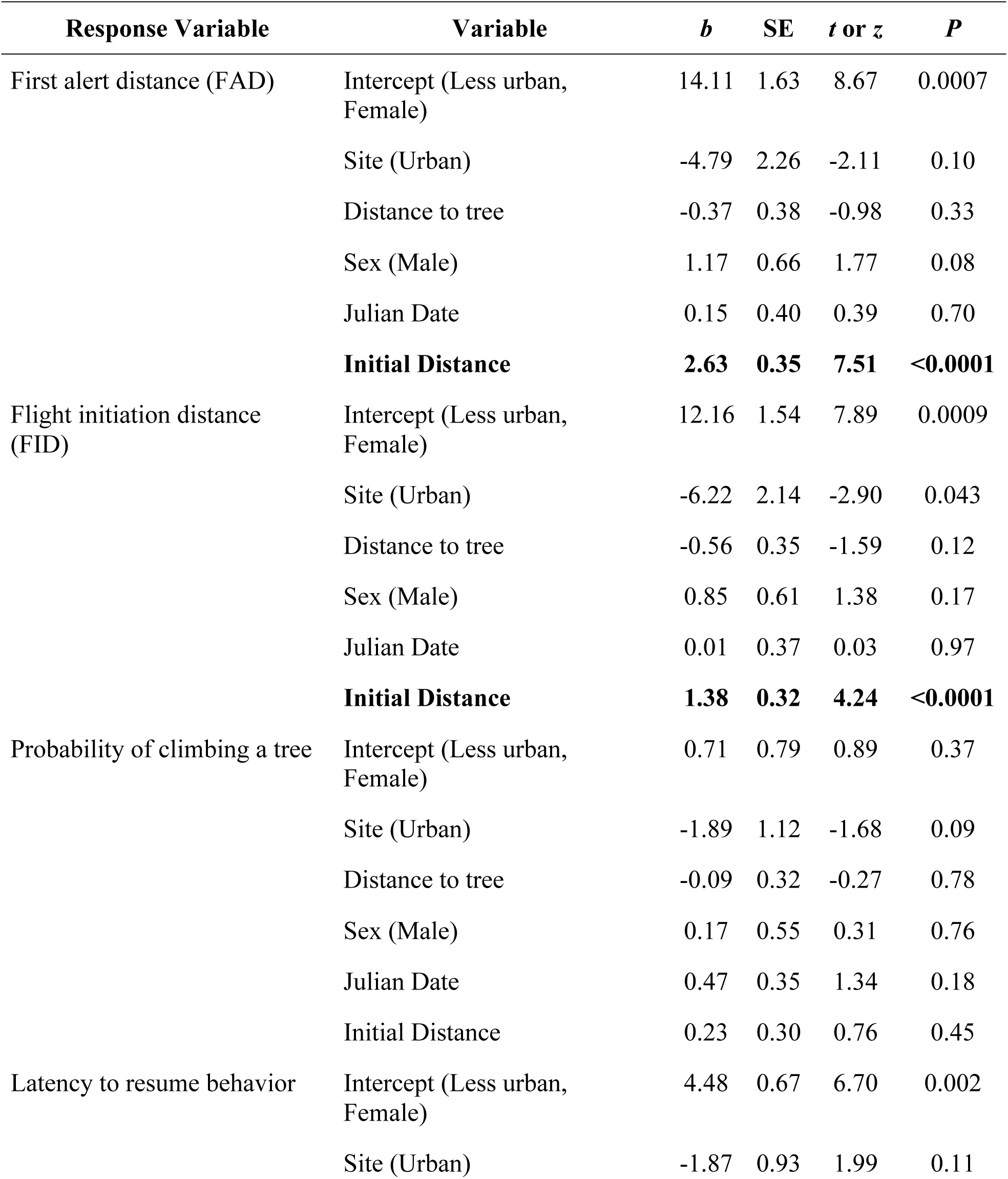
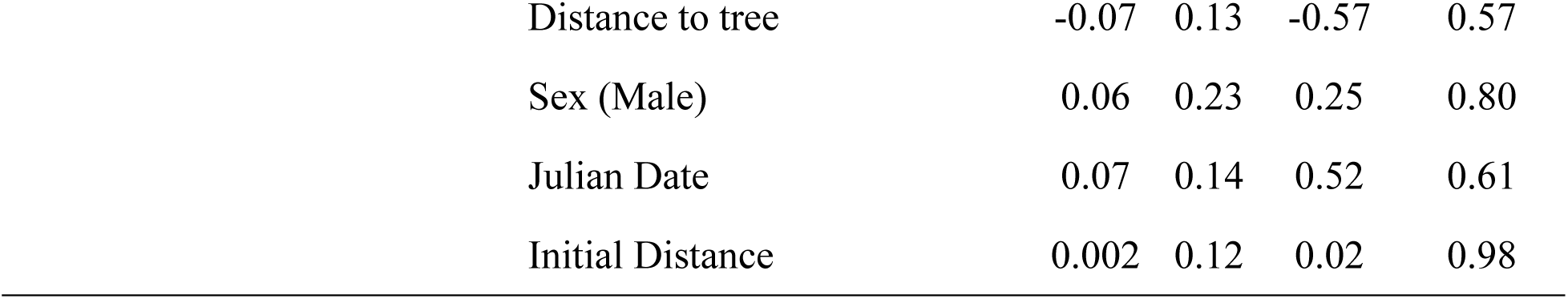
Differences between fox squirrels at urban and less urban sites that were not exposed to any playbacks for first alert distance (FAD), flight initiation distance (FID), probability of escaping the observer by climbing a tree during the trial, and latency to resume behavior following the trial. A random effect for site identity was included in the model or FAD (σ^2^ = 6.9), FID (σ^2^ = 6.24), probability of climbing a tree (σ^2^ = 1.23), and latency (σ^2^ = 1.22). Latency was ln+1 transformed. Results are from 77 trials from six sites.

| Response Variable | Variable | *b* | SE | *t* or *z* | *P* |
| --- | --- | --- | --- | --- | --- |
| First alert distance (FAD) | Intercept (Less urban, Female) | 14.11 | 1.63 | 8.67 | 0.0007 |
|  | Site (Urban) | -4.79 | 2.26 | -2.11 | 0.10 |
|  | Distance to tree | -0.37 | 0.38 | -0.98 | 0.33 |
|  | Sex (Male) | 1.17 | 0.66 | 1.77 | 0.08 |
|  | Julian Date | 0.15 | 0.40 | 0.39 | 0.70 |
|  | **Initial Distance** | **2.63** | **0.35** | **7.51** | **<0.0001** |
| Flight initiation distance (FID) | Intercept (Less urban, Female) | 12.16 | 1.54 | 7.89 | 0.0009 |
|  | Site (Urban) | -6.22 | 2.14 | -2.90 | 0.043 |
|  | Distance to tree | -0.56 | 0.35 | -1.59 | 0.12 |
|  | Sex (Male) | 0.85 | 0.61 | 1.38 | 0.17 |
|  | Julian Date | 0.01 | 0.37 | 0.03 | 0.97 |
|  | **Initial Distance** | **1.38** | **0.32** | **4.24** | **<0.0001** |
| Probability of climbing a tree | Intercept (Less urban, Female) | 0.71 | 0.79 | 0.89 | 0.37 |
|  | Site (Urban) | -1.89 | 1.12 | -1.68 | 0.09 |
|  | Distance to tree | -0.09 | 0.32 | -0.27 | 0.78 |
|  | Sex (Male) | 0.17 | 0.55 | 0.31 | 0.76 |
|  | Julian Date | 0.47 | 0.35 | 1.34 | 0.18 |
|  | Initial Distance | 0.23 | 0.30 | 0.76 | 0.45 |
| Latency to resume behavior | Intercept (Less urban, Female) | 4.48 | 0.67 | 6.70 | 0.002 |
|  | Site (Urban) | -1.87 | 0.93 | 1.99 | 0.11 |

|  |  |  |  |  |
| --- | --- | --- | --- | --- |
| Distance to tree | -0.07 | 0.13 | -0.57 | 0.57 |
| Sex (Male) | 0.06 | 0.23 | 0.25 | 0.80 |
| Julian Date | 0.07 | 0.14 | 0.52 | 0.61 |
| Initial Distance | 0.002 | 0.12 | 0.02 | 0.98 |

**Table 2.** Effects of acoustic playbacks (control, hawk, dog) on first alert distance (FAD), flight initiation distance (FID), probability of escaping human observer by climbing a tree, and latency to resume behavior following the trial for fox squirrels observed at urban or less urban sites. Reference (intercept) was “less urban” for site, “control playback” for treatment, and “female” for sex. A random effect for site identity was included in the model or FAD (σ^2^ = 6.5), FID (σ^2^ = 4.9), probability of climbing a tree (σ^2^ = 0.07), and latency (σ^2^ = 0.37). Latency was ln+1 transformed. Results are from 94 trials from six study sites.

| Response Variable | Variable | *b* | SE | *t* or *z* | *P* |
| --- | --- | --- | --- | --- | --- |
| First alert distance (FAD) | Intercept (Less urban, Female, Control) | 14.94 | 0.89 | 16.71 | <0.0001 |
|  | Site (Urban) | -2.32 | 1.33 | -1.74 | 0.11 |
|  | Dog playbacks | 0.51 | 0.90 | 0.57 | 0.57 |
|  | Hawk playbacks | -0.29 | 0.98 | -0.30 | 0.76 |
|  | Distance to tree | 0.36 | 0.29 | 1.23 | 0.22 |
|  | Sex (Male) | 0.29 | 0.58 | 0.51 | 0.61 |
|  | Julian Date | -0.10 | 0.28 | -0.34 | 0.73 |
|  | **Initial Distance** | **3.20** | **0.29** | **10.84** | **<0.0001** |
|  | Site (Urban) x Dog playbacks | 0.07 | 1.35 | 0.05 | 0.96 |
|  | **Site (Urban) x Hawk playbacks** | **4.29** | **1.49** | **2.86** | **0.005** |
| Flight initiation distance (FID) | Intercept (Less urban, Female, Control) | 12.3 | 0.95 | 7.31 | <0.0001 |
|  | Site (Urban) | -1.64 | 1.39 | -1.18 | 0.27 |
|  | **Dog playbacks** | **1.89** | **0.79** | **2.40** | **0.018** |
|  | Hawk playbacks | 1.21 | 0.85 | 1.42 | 0.16 |
|  | Distance to tree | 0.08 | 0.26 | 0.30 | 0.76 |
|  | Sex (Male) | 0.01 | 0.50 | 0.01 | 0.99 |
|  | Julian Date | -0.39 | 0.25 | -1.59 | 0.11 |
|  | **Initial Distance** | **2.99** | **0.26** | **11.51** | **<0.0001** |
|  | Site (Urban) x Dog playbacks | -1.52 | 1.18 | -1.29 | 0.20 |
|  | Site (Urban) x Hawk playbacks | 2.53 | 1.30 | 1.94 | 0.056 |
| Probability of climbing a tree | Intercept (Less urban, Female, Control) | -0.20 | 0.62 | -0.33 | 0.74 |
|  | Site (Urban) | -0.61 | 0.92 | -0.66 | 0.51 |
|  | Dog playbacks | 1.14 | 0.82 | 1.39 | 0.16 |
|  | Hawk playbacks | 1.26 | 0.90 | 1.40 | 0.16 |
|  | Distance to tree | -0.64 | 0.27 | -2.34 | 0.02 |
|  | Sex (Male) | 0.61 | 0.55 | 1.10 | 0.27 |
|  | Julian Date | -0.26 | 0.27 | -0.97 | 0.33 |
|  | Initial Distance | 0.01 | 0.27 | 0.04 | 0.96 |
|  | Site (Urban) x Dog playbacks | -1.52 | 1.20 | -1.27 | 0.20 |
|  | Site (Urban) x Hawk playbacks | 0.18 | 1.34 | 0.14 | 0.89 |
| Latency to resume behavior | Intercept (Less urban, Female, Control) | 3.83 | 0.39 | 9.69 | 0.0001 |
|  | Site (Urban) | -0.60 | 0.57 | -1.05 | 0.33 |
|  | **Dog playbacks** | **1.14** | **0.23** | **4.91** | **<0.0001** |
|  | **Hawk playbacks** | **1.71** | **0.25** | **6.76** | **<0.0001** |
|  | Distance to tree | 0.03 | 0.07 | 0.39 | 0.70 |
|  | Sex (Male) | 0.24 | 0.15 | 1.61 | 0.11 |
|  | Julian Date | 0.07 | 0.07 | -0.91 | 0.36 |
|  | Initial Distance | 0.06 | 0.08 | 0.82 | 0.41 |
|  | Site (Urban) x Dog playbacks | 0.03 | 0.35 | 0.10 | 0.92 |
|  | Site (Urban) x Hawk playbacks | 0.22 | 0.39 | 0.57 | 0.57 |

### Quantifying behavioral responses of squirrels

In total, we observed fox squirrels for 52.36 hours over 30 different days. A single observer (AK) conducted all trials. A total of 171 trials were conducted with 71 trials conducted without any acoustic playback treatments and 94 trials conducted with an acoustic playback. Sites were visited between 800 and 1900 h EST and data were collected from October 2019 through January 2020. Trials with no playbacks were conducted from 25 October 2019 to 16 November 2019 (from 812 to 1810 h) whereas trials using playbacks (playback trials) were conducted from 18 November 2019 to 27 January 2020 (from 802 to 1609 h). We randomized the order in which sites were visited and the version of playback treatments used (see below for information on playback versions) at each site. No two sites were visited on the same day. All trials with no playbacks were conducted prior to the playback trials in this study. This was due to personnel limitations and prevents us from directly comparing trials with and without playbacks given that squirrel behavior likely changes seasonally from October to January due to food caching behavior in autumn but not winter. Trials were not conducted when it was raining or snowing. Results from two one-way ANOVAs showed that the time of day for playback trials did not vary among the three different treatment groups (*F*_2,91_ = 0.44, *p* = 0.65) and that the time of day when trials were conducted did not vary among squirrels at the urban or less urban sites (t-test: *t*_75.4_ = 0.21, *p* = 0.83). Air temperature varied during all the trials varied from -6.1° to 10.5°C (mean = 2.7° C).

We measured the behavioral responses of squirrels to humans or humans plus different playbacks using protocols developed for tree squirrels (Dill and Houtman, 1989; Gustafson and VanDruff, 1990; McCleery, 2009). At the beginning of each trial, a marker was placed at the starting position of the observer and trial data were recorded (GPS location, time of day, temperature, general weather conditions, and squirrel sex). The squirrel was approached by a single observer (AK) at a slow and steady pace in a direct line to the squirrel (see Fig. S1 in Appendix I). Additional markers were placed when a squirrel displayed the first alert and when they fled. The FAD was defined as the distance between the observer and the squirrel when it first stopped moving (froze) and looked at the observer with one or both eyes. FID was recorded as the distance between the squirrel’s initial position and the observer location when it actively fled (stopped feeding and foraging and moved rapidly away from observer). A marker was also placed at an estimate of the squirrel’s initial position to the place where they fled to if refuge was not taken in a tree, and the distance between this marker and the squirrel’s initial position was recorded as “flight distance”. We recorded this because some studies have noted that FID is variable depending on intruder starting distances and distance to a refuge (Dill & Houtman, 1989; Blumstein, 2003). Consequently, the distance between the observer and the focal squirrel at the start of the trial (starting positions, hereafter referred to as “initial distance”: mean ± SE over all 171 trials = 21.2±0.61 m) and the distance between the focal squirrel’s initial position and the nearest tree were also measured (“distance to nearest tree”: mean ± SE over all 171 trials =3.2±0.11 m).

If the focal animal took refuge in a tree other than the one nearest, distance between the squirrel’s initial position and its refuge tree of choice (“distance to the chosen tree”) was also recorded. If an individual took refuge in a tree, a laser rangefinder was used to measure how high they climbed, and this distance was recorded (same as vertical escape distance in Uchida et al., 2017). Lastly, latency to resume behavior was recorded (“latency”). This was measured with a stopwatch to determine how long it took for the animal to cease alert/vigilance behavior and resume typical activity (foraging or traveling off tree). When the observer was recording latency, they maintained as large a distance as possible (∼15-20 m) from the tree to reduce the influence on the squirrel’s behavior. Out of all the trials, nearly all squirrels ceased alert behavior and resumed typical behavior within a couple minutes (n = 171 trials, mean ± SE = 130.5 ± 10.3 s), but there was one individual squirrel that remained alert for longer than ten minutes and we recorded its latency as ten minutes. Distances were measured with a tape measure and presented in meters.

### Playback trials

Playback trials (n = 94 total trials) were conducted using the same protocol shown above, with the addition of an acoustic stimulus being broadcasted while the observer approached the squirrels. The control stimulus consisted of recordings of black-capped chickadee calls (*Poecile atricapillus*). Black-capped chickadees are not known to be predators of fox squirrels (Korschgen, 1981; Koprowski, 1994) and a previous study in another tree squirrel species showed that individuals exhibited a significantly reduced response to black-capped chickadee playbacks compared to calls of other anthropogenic noises (car alarm, buzzer) and playbacks of red-tailed hawks (Bohls and Koehnle, 2017). We therefore expected that black-capped chickadee recordings would represent a neutral vocalization for fox squirrels and they can act as control to ensure that any differences in squirrel behavior across playback treatments are attributable to the vocalization information of the playback rather than an added exposure to noise. To simulate the threat of a terrestrial predator, recordings of domestic dogs barking were broadcasted. Domestic dogs are terrestrial predators of fox squirrels (Koprowski, 1994; Wauters et al., 1997) and other species of tree squirrels that live in the same habitats as fox squirrels also adjust their risk-taking behavior according to the abundance of domestic dogs (Bowers and Breland, 1996; Cooper et al., 2008). For the avian predator, recordings of red-tailed hawk (*Buteo jamaicensis*) calls were broadcasted to the focal individual. Red-tailed hawks were chosen since they are year-round predators of fox squirrels in Michigan (Koprowski, 1994; personal observations) and other studies illustrate that tree squirrels respond to hawk playbacks with increased anti-predator behavior (McCleery, 2009; Lilly et al., 2019). No post-processing of sound files was performed.

Playbacks of the recordings were broadcasted to individuals at the start of the trial and when the observer began the approach and suspended when the squirrel took flight. Vocalizations were broadcasted through a JAMBOX speaker (Jawbone, San Francisco, CA) connected to an Apple iPhone 6s (Mountain View, CA) with a constant volume set for the speaker and phone. The speaker was carried by the observer during each trial. The amplitude of the playbacks measured from 1 m away from the speaker was variable among the chickadee (67-80 dB), dog (69-77 dB), and hawk (78-86 dB) playbacks (measured using a BAFX Sound Level Meter, BAFX3370). We note that the initial starting distance of the playbacks was inherently variable as we could not standardize the distance between the observer and squirrel when the trials were started (mean ± SE over all 94 trials involving playbacks = 19.11±0.73 m). Consequently, the actual realized sound level of the playbacks experienced by a squirrel varied. Given how the trials were conducted in real time (not video recorded), the single observer (AK) was not blind to the playback treatments or locations of where the experiments took place. All vocalization recordings were found online (Control A: Place, 2015; Control B: Floyd, 2017a; Control C: Floyd, 2017b; Dog A: Simion, 2016; Dog B: Simion, 2018; Dog C: Simion, 2017; Hawk A: Chartier, 2008; Hawk B: Addison, 2017; Hawk C: Wilson, 2010). Each playback treatment (control, dog, or hawk) had three separate recordings/exemplars (A, B, or C). We tested whether there were any exemplar effects in separate ANOVAs that included playback exemplar (A, B, C), playback treatment (chickadee, dog, hawk), and an interaction between the two. We did this for each of our four behavioral response variables and did not find any significant interactions between playback exemplar and treatment (*p* = 0.12-0.99), suggesting that the version of the playback treatment did not influence the behavioral response.

### Statistical analyses

We analyzed the data from trials with and without playbacks separately because the two experiments were not conducted synchronously and seasonal changes from fall to winter in Michigan may alter squirrel behavior. In trials without playbacks, we used three separate linear mixed-effects models (LMMs) to examine the effects of urbanization on FAD, FID, and latency to resume activity following the trial. Although the linear distance a squirrel climbed up a tree (from base of tree to location of squirrel) has been used in other studies of tree squirrels (e.g., Uchida et al., 2017), the distance a squirrel climbed up a tree in our study exhibited a Poisson distribution where many squirrels did not climb up a tree at all and a few climbed up very high (squirrels did not climb a tree in 80 of 171 total trials; those that did climb a tree mean ± SE = 4.7 ± 0.34 m). This seemed to better approximate a behavioral decision made by a squirrel to “climb or not climb” rather than “how high to climb”. Consequently, a generalized linear mixed-effect model (GLMM) with binomial errors was used to examine the effects of urbanization on the probability that squirrels climbed a tree to escape during the trial. We note that the same inferences for the linear distance a squirrel climbed a tree were gained if we instead ran a zero-inflated Poisson mixed-effects model. Models included site category (urban, less urban), distance to the nearest tree, sex, Julian date of the trial, and initial distance of the observer as fixed effects. Distance to the nearest tree was included not only because previous studies show it can impact anti-predator behavior (measured using FID: Dill & Houtman, 1989; Blumstein, 2003) but also because it helps control for any differences in vegetation among the different study sites, which could impact their behavior. Because we had repeated samples from the same site, we also included a random intercept for site in all of our models. The same model structure was used in separate LMMs or the GLMM for data from the playback trials to examine the effects of the acoustic playback manipulations on the four squirrel behaviors described above but the models included an interaction between playback treatment (control, dog, hawk) and site category (urban, less urban). We then assessed the statistical significance of pairwise comparisons using post-hoc Tukey’s Honest Significant Differences that were corrected for multiple comparisons. In these pairwise comparisons, we were specifically interested in identifying 1) whether squirrels in urban and less urban sites differed in their behavior in response to the playback treatments (e.g., urban squirrels exposed to hawk playbacks differed in FID compared to less urban squirrels exposed to hawk playbacks) and 2) whether squirrels within each type of site differed in their response to the playbacks (e.g., whether squirrels in urban areas exhibited a higher FID in response to hawk playbacks compared to those in urban areas exposed to control playbacks).

Continuous predictor variables were standardized to a mean of 0 and SD of 1. We confirmed model diagnostics visually and all models met the appropriate assumptions (normality of residuals, constant variance, no high leverage observations). Latency to resume behavior was log+1 transformed (base e) to improve homoscedasticity and normality. There were also no predictor variables that were found to be colinear as all variance inflation factors (VIFs) were less than 3.68 (Zuur et al., 2010), though the higher VIF were due to interaction terms and VIF of variables not in interactions were <1.5. All analyses were conducted in R version 4.02 (R Core Team, 2020) with lme4 (version 1.1.23, Bates et al., 2015) and p-values estimated using lmerTest (version 3.1.2, Kuznetsova et al., 2017). Tukey’s post-hoc tests were used to evaluate if the responses to the playback treatments differed between squirrels in urban and less urban areas using emmeans (1.5.2-1: Lenth, 2020) and *p* values from these analyses were adjusted for multiple comparisons. Mean and SE are presented below.

## Results

### Behavioral responses to human-stimuli only

Overall, urban squirrels (n = 38, 20 females and 18 males) exhibited greater tolerance to humans as they allowed a human observer to get closer to them before they exhibited vigilance (FAD) or fled (FID) and tended to be less likely to climb a tree during the trial and more quickly return to typical behavior following the trial compared to those in less urban sites (n = 39, 19 females and 20 males; Table 1, Fig. 1). FID in the squirrels at the urban site (6.36 ± 0.52 m) was 97.2% shorter compared to those at the less urban sites (12.54 ± 0.62 m, *p* = 0.043, Table 1, Fig. 1B). Although the average FAD for squirrels observed at the urban site (10.09 ± 0.74 m) was 43.8% shorter than for those at the less urban sites (14.51 ± 0.69 m), this difference was not significant (*p* = 0.10, Table 1, Fig. 1A). Squirrels at the urban sites were less likely to climb a tree while the observer approached (34.2% of trials) compared to those at the less urban sites (64.1%), although this difference was not significant (*p* = 0.093, Table 1, Fig. 1C). Latency to resume behavior following the trial was shorter for urban squirrels (43.9 ± 14.4 s) compared to those at the less urban sites (157.51 ± 25.35 s), but this difference was not significant (*p* = 0.11, Table 1, Fig. 1D). Trials where the observer started the trial at a longer initial distance to the squirrel had significantly longer FAD and FID but not probability of climbing a tree or latency to resume behavior following the trial (Table 1). There were no significant effects of sex, Julian date, or distance to the nearest tree on FAD, FID, probability of climbing a tree, or latency (Table 1).

**Figure 1.**
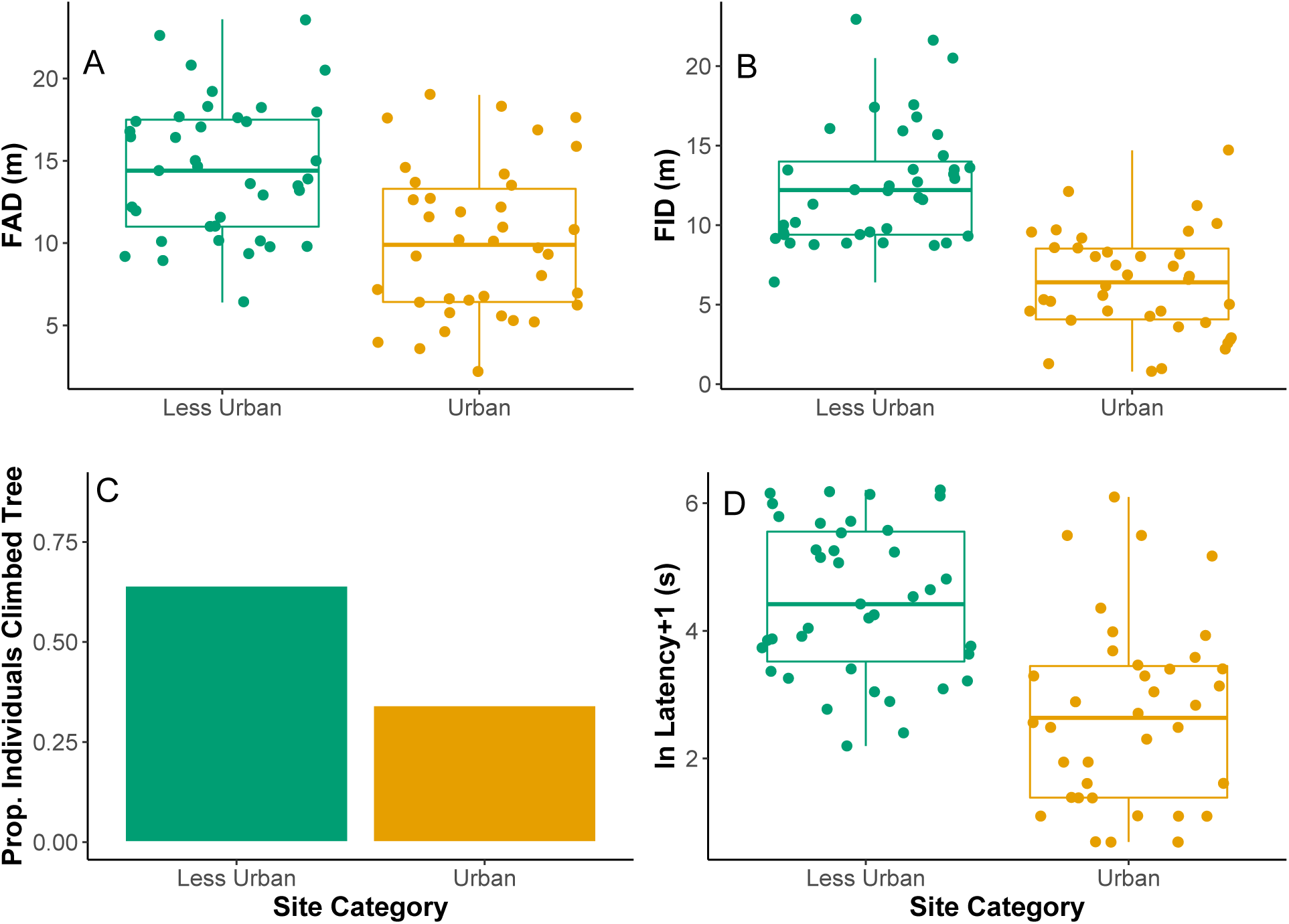
Variation in A) vigilance (first alert distance: FAD), B) anti-predator behavior (flight initiation distance: FID), C) proportion of individuals that escaped up a tree during the trial, and D) latency to resume behavior following the trial among fox squirrels at urban (n = 38 trials) and less urban (n = 39) sites in trials where squirrels were not exposed to any playbacks. Squirrels in urban areas had significantly shorter FID compared to those in the less urban sites, but there were no other significant differences (Table 1). Each symbol corresponds to a different trial. Upper and lower hinges correspond to first and third quartile, respectively. Upper/lower whiskers extend from the hinge to the highest/lowest value that is within 1.5x the interquartile range. Solid horizontal line shows median.

### Behavioral responses to stimuli from natural predators

The effects of the playbacks on FAD depended upon whether the squirrels were located at the urban or less urban sites (Table 2, Fig. 2A). Average FAD for urban squirrels exposed to the hawk vocalizations (n = 12 trials, 17.17 ± 1.33 m) was 37% longer than urban squirrels who were exposed to the control playback (n = 11, 12.56 ± 0.94 m, Tukey’s *p* = 0.004) and 20% longer than those exposed to the dog playbacks (n = 20, 14.30 ± 0.83 m, Tukey’s *p* = 0.006, Fig. 2A). By contrast, the FAD of squirrels at the less urban sites were just longer overall (Fig. 2A) and the FAD of those less urban squirrels who were exposed to the hawk playbacks (n = 16, 14.62 ± 1.14 m) did not differ from those exposed to the control playback (n = 15, 14.46 ± 0.88 m, Tukey’s *p* = 0.99) or dog playbacks (n = 20, 14.85 ± 1.05 m, Tukey’s *p* = 0.94, Fig. 2A). There were no significant differences in FAD for squirrels exposed to dog playbacks and those exposed to the control playback for squirrels at urban sites (Tukey’s *p* = 0.99) or those at less urban sites (Tukey’s *p* = 0.99). When comparing squirrels at urban or less urban sites to a specific playback treatment, squirrels at the urban and less urban sites did not differ in their FAD when exposed to hawk playbacks (urban vs. less urban: Tukey’s *p* = 0.64), dog playbacks (urban vs. less urban: Tukey’s *p* = 0.39), or the control stimulus (urban vs. less urban: Tukey’s *p* = 0.49, Fig. 2A).

**Figure 2.**
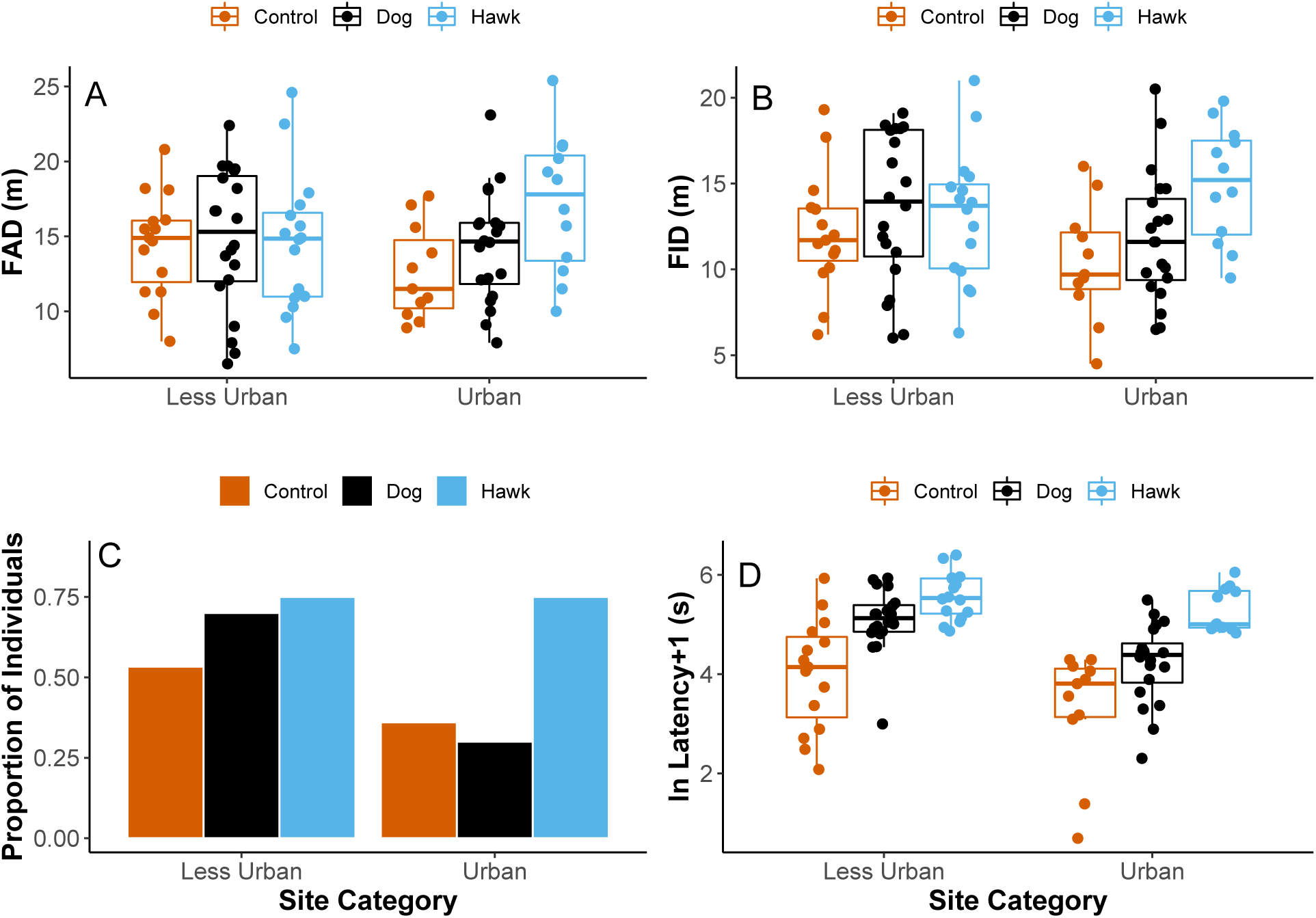
Effects of human observer approaching a squirrel while broadcasting one of three playback treatments (control playback, hawk or dog vocalizations) on A) first alert distance (FAD), B) flight initiation distance (FID), C) proportion of individuals escaping up a tree during the trial, and D) latency to resume typical behavior following the trial. Trials were conducted at less urban (n = 15 control, 20 dog, 16 hawk) and urban (n = 11 control, 20 dog, 12 hawk) sites. Results shown in Table 2. Upper and lower hinges correspond to first and third quartile, respectively. Each symbol corresponds to a different trial. Upper/lower whiskers extend from the hinge to the highest/lowest value that is within 1.5x the interquartile range. Solid horizontal line shows median.

Similar to FAD, the effects of the playbacks on FID also depended upon whether the squirrels were located at the urban or less urban sites (Table 2, Fig. 2B). Average FID for urban squirrels exposed to the hawk vocalizations (n = 12 trials, 14.95 ± 0.98 m) was 44.3% longer than squirrels who were exposed to the control playbacks (n = 11, 10.37 ± 3.13 m, Tukey’s *p* = 0.001) and 29.9% longer than those exposed to the dog playbacks (n = 20, 11.86 ± 2.65 m, Tukey’s *p* = 0.001, Fig. 2A). In squirrels at urban sites, there was no difference in FID between those exposed to the control playback and those exposed to the dog vocalizations (Tukey’s *p* = 0.99). In squirrels at less urban sites, FID for those exposed to the dog playbacks (13.60 ± 0.98 m) was similar to those exposed to the control (12.12 ± 0.88 m, Tukey’s *p* = 0.14) or hawk playbacks (13.11 ± 0.96 m, Tukey’s *p* = 0.95, Fig. 2B). Unlike urban squirrels, the FID of those at the less urban sites who were exposed to hawk playbacks was similar compared to those exposed to the control playback (Tukey’s *p* = 0.69). When comparing squirrels at urban or less urban sites to a specific playback treatment, squirrels at the urban and less urban sites did not differ in their FID when exposed to hawk playbacks (urban vs. less urban: Tukey’s *p* = 0.98), dog playbacks (urban vs. less urban: Tukey’s *p* = 0.13), or the control stimulus (urban vs. less urban: Tukey’s *p* = 0.83).

There were no significant effects of the playback treatments or site differences on the likelihood squirrels climbed a tree. Squirrels at urban sites were not more likely to climb a tree when exposed to a hawk playback compared to a dog (Tukey’s *p* = 0.30) or control (Tukey’s *p* = 0.67) playback and were not more likely to climb a tree when exposed to a dog playback versus a control playback (Tukey’s *p* = 0.99). Squirrels at less urban sites exhibited a similar probability of climbing a tree when they were exposed to hawk playbacks compared to dog (Tukey’s *p* = 1.0) or control (Tukey’s *p* = 0.73) playbacks or when exposed to dog playbacks compared to a control playback (Tukey’s *p* = 0.73). Squirrels at the urban and less urban sites did not differ in their probability of climbing a tree when exposed to hawk playbacks (urban vs. less urban: Tukey’s *p* = 0.99), dog playbacks (urban vs. less urban: Tukey’s *p* = 0.12), or the control stimulus (urban vs. less urban: Tukey’s *p* = 0.99).

Squirrels at the urban and less urban sites did not differ in their latency to resume typical behavior following exposure to hawk playbacks (urban vs. less urban: Tukey’s *p* = 0.98), dog playbacks (urban vs. less urban: Tukey’s *p* = 0.89), or the control stimulus (urban vs. less urban: Tukey’s *p* = 0.88, Table 2, Fig. 2D). However, there were differences in how squirrels responded to the playback treatments within each of the two types of study sites. Squirrels at urban sites that were exposed to the hawk playbacks took 436% longer to resume their pre-trial behavior (213.33 ± 28.9 s) compared to those who were exposed to the control playback (39.82 ± 7.70 s, Tukey’s *p* < 0.001) and 147% longer than those exposed to dog playbacks (86.25 ± 13.30 s, Tukey’s *p* = 0.027). Squirrels at the urban sites also took 114% longer to resume typical behavior if they were exposed to dog playbacks compared to those exposed to the control playback (Tukey’s *p* < 0.001). Similarly, squirrels at the less urban sites that were exposed to the hawk playbacks took 218% longer to resume their pre-trial behavior (291.25 ± 35.45 s) compared to those who were exposed to the control playback (91.47 ± 25.45 s, Tukey’s *p* < 0.001) and 55.7% longer than those exposed to dog playbacks (187.05 ± 21.38 s, Tukey’s *p* = 0.12, Fig. 2D). Squirrels at the less urban sites also took 95.6% longer to resume typical behavior if they were exposed to dog playbacks compared to those exposed to the control playback (Tukey’s *p* < 0.001).

There was no effect of sex or Julian date of trial, on any of the behaviors (Table 2). There was no effect of the initial distance that a squirrel was from a tree when the trial started on FAD, FID, or probability to climb a tree, but squirrels were less likely to climb a tree if they were closer to one when the trial started (Table 2). Trials that started with the human observer a greater distance away from the squirrel had longer FAD, FID, and latency to resume typical behavior, but not the probability to climb a tree (Table 2).

## Discussion

Squirrels at urban sites in the no playback trials exhibited a significantly shorter FID compared to those at the less urban sites and also exhibited a lower FAD and likelihood to climb a tree during the trial, and shorter latency to resume typical behavior following the trial, though only the difference in FID was statistically significant. In the trials where squirrels were exposed to playbacks from possible predators (hawks or dogs), squirrels at the urban sites did not differ in their vigilance (FAD) or anti-predator behavior response (FID, likelihood to climb a tree, latency to resume typical behavior following the trial) compared to those at the less urban sites. When we compared the behavior responses of squirrels within each site type (urban or less urban), squirrels at the urban sites exhibited longer FAD (hawk > dog = control) and FID (hawk > dog = control) when exposed to hawk playbacks compared to control or dog playbacks, suggesting increased vigilance (FAD) anti-predator behavior (FID) when exposed to vocalizations from potential predators. By contrast, squirrels at the less urban sites had longer overall FAD and FID than those at urban sites regardless of playback treatment and there was no effect of hawk or dog playbacks on FAD (hawk = control = dog) or FID (hawk = control = dog), suggesting no increase in vigilance or anti-predator behavior when exposed to vocalizations from potential predators. Squirrels at both urban and less urban sites were not more likely to climb a tree following playbacks from possible predators (hawk = dog = control) but both urban and less urban squirrels exhibited a longer latency to resume typical behavior following the hawk or dog playbacks compared to the control (hawk > dog > control), suggesting increased anti-predator behavior when exposed to vocalizations from potential predators. Overall, our results indicate that squirrels in urban areas are more tolerant to humans but still exhibit a high level of vigilance and anti-predator behavior when exposed to predator stimuli. In terms of the STRANGEness of our results (Webster and Rutz, 2020), our results may be generalizable to other squirrel populations or different species but we note that our results are biased towards squirrels that voluntarily participated in the trials (i.e., did not run away when approached). We also note that the significance of our results may be limited to urban populations where predators are present in those areas.

Similar to most other studies in terrestrial animals (Samia et al., 2015) and in studies in tree squirrels (McCleery, 2009; Engelhardt and Weladji, 2011; Sarno et al., 2015; Uchida et al., 2020), our results from trials with no playbacks suggest that squirrels in urban sites were more tolerant of humans. Specifically, squirrels in urban areas exhibited a shorter FAD and FID, lower probability to climb a tree to escape the human observer, and a shorter latency to resume typical behavior following the trial, although only FID was significantly different between habitat types. The congruency of our results with previous studies strongly supports this assumption that squirrels at our urban sites were more tolerant of humans. These are presumably sympatric populations with a large amount of gene flow among them as the linear distance between some urban and less urban sites is ∼1 km. Unless selection favoring reductions in anti-predator behavior is extremely strong in urban areas or features of urban landscapes strongly impede gene flow (Johnson and Munshi-South, 2017), it seems likely that these behavioral differences are driven by plasticity given that the likely exchange of individuals between suburban and urban sites prevents local genetic adaptation to these different sites (see discussion in Sol et al., 2013). It is also possible that these behavioral differences are due to personality-dependent colonization of urban habitats (Carrete and Tella, 2010; Sprau and Dingemanse, 2017), but we cannot distinguish among these possibilities at this time.

Although squirrels at our urban sites were more tolerant of humans, they still exhibited a strong behavioral response to acoustic stimuli from natural predators. Specifically, they exhibited increased vigilance (FAD) and anti-predator behavior (FID, latency to resume typical behavior after the trial) when exposed to the playbacks of a natural predator (hawk) compared to the control playback or the dog playbacks. The behavioral responsiveness to hawk vocalization is somewhat surprising because hawks do not vocalize while hunting, but squirrels still responded to their presence suggested through acoustic cues. These results indicate that urban squirrels do still pay attention to predation risk and can discriminate and respond accordingly by becoming vigilant and fleeing when the human is at a greater distance if the human is also paired with hawk playbacks. By contrast, squirrels at less urban sites did not exhibit differences in FAD when exposed to the different playbacks, perhaps due to some ceiling effect given that FAD of squirrels at less urban sites was much longer than FAD of squirrels at urban sites. Furthermore, when we compared the effects of hawk or dog playbacks on FAD or FID, there were no differences between squirrels at the urban and less urban sites. Our results therefore reject the hypothesis that urban squirrels are less responsive to natural predators due to increased tolerance to humans. Previous studies (see also Labra and Leonard, 1999; Seress et al., 2011; Cooper et al., 2008; Bokony et al., 2012; Cavalli et al., 2016; Weaver et al., 2018; Vincze et al. 2019) together with our results support that animals in urban habitats or those frequently exposed to humans, even if more tolerant of human presence, still exhibit increases in anti-predator behavior in response to a non-human predator. However, future studies that test this hypothesis need to have increased sample sizes and should also include a playback treatment that uses both visual and acoustic cues of humans as a control stimulus.

There are two other interesting results from our trials with playbacks. First is the finding that squirrels that were closer to a tree at the start of the trial were less likely to climb a tree. This is opposite of what we would expect and future studies need to better assess if this is caused by some larger habitat difference between urban and less urban areas and/or reflect differential escape strategies. For example, squirrels at urban sites may be more distant to a tree at the start of the trials and escape from humans by running away rather than going up a tree. However, in a post-hoc analysis using our entire dataset of trials conducted with or without playbacks (n = 171 trials), squirrels in urban areas (n = 81 trials, 3.30 ± 0.16 m) and those in less urban areas (n = 90 trials, 3.05 ± 0.15 m) did not differ in their distance to a tree at the start of the trials (general linear model: *t*_169_ = 1.13, *p* = 0.26). Second, we expected that squirrels would respond to the dog playbacks in similar way to how they responded to the hawk playbacks as both are stimuli from potential predators. Previous studies in tree squirrels also show that they exhibit increased vigilance or FID to the physical presence of a dog with a human handler (Gustafson and VanDruff, 1990; Cooper et al., 2008) or behave in such a way in areas with high levels of dogs and cats that suggests that they perceive a higher predation risk in such areas (i.e., giving up density was higher in study areas where cats and dogs are present: Bowers and Breland, 1996). Instead, we found that squirrels at both sites did not differ in their vigilance (FAD) when exposed to dog playbacks compared to the control playbacks. Additionally, only squirrels in less urban sites had a slightly (non-significantly) longer FID when exposed to dog playbacks compared to the control playback. Although we did find that squirrels exposed to dog vocalizations took longer to resume typical behavior following the trials, our results generally differ from previous studies in tree squirrels and owls showing that FAD and/or FID were increased in squirrels or owls in urban areas when they were presented with a human plus dog compared to just a human (Gustafson and VanDruff, 1990; Cooper et al., 2008; Cavalli et al., 2016). This suggests that squirrels, especially those in urban sites, were more tolerant to dog vocalizations when paired with a human observer, whereas squirrels in less urban areas may have viewed the sounds of dogs paired with humans as threatening. We predict that squirrels in urban environments exhibit selective tolerance where the response to humans or stimuli from their commensals (dogs) is attenuated but the increased response to natural predators is maintained despite this tolerance to humans and their dogs.

Together, our results provide insight into how urbanization may shape the behavioral characteristics animals in two main ways. First, as most studies on this topic are in birds (Samia et al., 2015), which can escape from humans using flight, it is important to consider if the same patterns are found in terrestrial animals. Our results show that non-volant animals in urban environments exhibit less vigilance and anti-predator behavior. Second, we show that squirrels in urban environments were more tolerant to humans but still exhibited a strong response reflecting increased vigilance and anti-predator behavior to acoustic stimuli from a natural predator (hawks) and that squirrels in urban areas did not differ in their behavioral response when exposed to stimuli from two types of possible predators compared to those exposed to those stimuli in less urban areas. Although we do not wish to imply that tolerance or habituation to humans is cost-free, most studies fail to find evidence that populations where individuals are more tolerant of humans (or in some cases habituated to their presence) also exhibit reduced vigilance and/or anti-predator behavior to stimuli from natural predators. Given that urbanization is unlikely to slow, increased effort is needed to determine if increased tolerance and/or habituation to humans carries costs (Geffroy et al., 2015). Some studies suggest the costs of human tolerance may be more nuanced, such as tolerance to humans reducing the latency to return to the nest following a disturbance in nesting shorebirds, but potentially causing increased chick mortality due to the presence of dogs that often are paired with human stimuli (Baudains and Lloyd, 2007). Other studies that increased tolerance of humans could even be beneficial for populations that cannot avoid anthropogenic stimuli due to seasonal food pulses coinciding with a large influx of tourists (Wheat and Wilmers, 2016). Clearly more work is needed on this subject, especially on a greater number of species including species that are not “urban exploiters” like tree squirrels, but the existing evidence rejects the hypothesis that there is a cost to human tolerance in terms of lowering the vigilance and/or anti-predator behavior of animals to other natural predators. If predatory species re-colonize urban areas, our results suggest that they should respond appropriately to stimuli indicating their presence.

## Data Availability Statement

All data are available through the FigShare account associated with the senior author (Dantzer, 2021).

## Acknowledgements

This research was a part of a senior honor’s thesis conducted by AK in the Department of Psychology at the University of Michigan. It was funded by the University of Michigan. Thanks to Amy-Charlotte Devitz for input on this project and to two anonymous reviewers for constructive feedback. We acknowledge that the University of Michigan and some of the other properties that we worked on during this project reside on the traditional Territories of the Three Fire Peoples – the Ojibwe, Odawa, and Bodewadmi.

